# Egocentric Biases are Determined by the Precision of Self-related Predictions

**DOI:** 10.1101/2021.04.02.437869

**Authors:** Leora Sevi, Mirta Stantic, Jennifer Murphy, Michel-Pierre Coll, Caroline Catmur, Geoffrey Bird

## Abstract

According to predictive processing theories, emotional inference involves simultaneously minimising discrepancies between predictions and sensory data relating to both one’s own and others’ states, achievable by altering either one’s own state (empathy) or perception of another’s state (egocentric bias) so they are more congruent. We tested a key hypothesis of these accounts, that predictions are weighted in inference according to their precision (inverse variance). If correct, more precise self-related predictions should bias perception of another’s emotional expression to a greater extent than less precise predictions. We manipulated predictions about upcoming own-pain (low or high magnitude) using cues that afforded either precise (a narrow range of possible magnitudes) or imprecise (a wide range) predictions. Participants judged pained facial expressions presented concurrently with own-pain to be more intense when own-pain was greater, and precise cues increased this biasing effect. Implications of conceptualising interpersonal influence in terms of predictive processing are discussed.

---

The notion that the brain is an inferential machine, generating predictions to explain the sensory data it receives in order to test models about the state of the world, is becoming increasingly influential in cognitive neuroscience. Within the predictive processing framework (Clark, 2016; Friston, 2010; Hohwy, 2013), the brain continually tests predictions about the world, generated by models, against incoming sensory data. Discrepancies between predictions and sensory data (prediction errors) are resolved either through, 1) the updating of models generating predictions such that they better fit sensory data, or 2) the performance of ‘action’ (whether cognitive, motoric or interoceptive), in order to minimise the discrepancy between predictions and sensory data. Which of these strategies are enacted in order to reduce prediction errors is a function of the relative expected precision (uncertainty, confidence — or, mathematically, inverse variance) of predictions and prediction errors: if prediction errors are more precise than predictions then models are updated; if not, action occurs.

One feature of the models specified by predictive processing theories is that they are hierarchical; at lower levels, they attempt to explain unimodal sensory data, whereas at higher levels they are multimodal, generating exteroceptive (e.g., visual, auditory), proprioceptive, and interoceptive (e.g., hunger, satiety, pain) predictions. These higher levels allow predictions and prediction errors relating to contingent events in different modalities to contextualise each other, allowing for more abstract representations of a cause of sensory data, including the action goals (Kilner, Friston, & Frith, 2007), mental states (Friston & Frith, 2015; Koster-Hale & Saxe, 2013) and affective states (Barrett & Simmons, 2015; Demekas, Parr, & Friston, 2020; Ondobaka, Kilner, & Friston, 2017; Peng, Huang, Liu, & Cui, 2019; Quattrocki & Friston, 2014; Seth, 2013; Seth & Friston, 2016) of ourselves and other people. Importantly, this feature allows action in one modality to resolve prediction error in another (Pezzulo, Rigoli, & Friston, 2015), across individuals. Thus, models can link exteroceptive predictions about states of the other and interoceptive/proprioceptive predictions about the states of the self, and should either of these fail to explain all the sensory data, then prediction error in one domain can be reduced via ‘action’ in another. This means that exteroceptive predictions concerning states of the other can induce change in the states of the self via interoceptive/proprioceptive ‘action’, and interoceptive/proprioceptive predictions concerning states of the self can induce change in the perception of another’s state via exteroceptive ‘action’.

As an example, consider the case in which one agent, Derek, observes another agent, Rodney, in pain. In order to estimate the causes of the exteroceptive sensory data before him (i.e., Rodney’s pained expression), Derek can use his own model of pain. Providing Derek has experienced a developmental environment in which others (e.g., caregivers) responded to his pain by displaying pained expressions/vocalisations themselves, Derek’s pain model will include predictions relating both to the sight/sound of another in pain and the feeling of pain in himself (Bird & Viding, 2014; Heyes & Bird, 2007). Thus, activation of Derek’s pain model will generate both interoceptive (what pain will feel like) and exteroceptive (e.g., what another’s face will look like) predictions. These exteroceptive predictions will provide a good fit to the exteroceptive sensory data (Rodney’s pained expression). However, the interoceptive predictions about Derek’s own pain would not be fulfilled in this situation and so prediction errors would be generated. As outlined earlier, these prediction errors could be resolved if the predictions cause the instantiation of a pained state in Derek (interoceptive action), i.e., they cause Derek to feel empathy for Rodney.

Meanwhile, Rodney’s pain model, if it is the same as Derek’s, is generating the same interoceptive and exteroceptive predictions. The interoceptive predictions are fulfilled by Rodney’s own pain and, if Derek did indeed empathise with Rodney and make a pained expression, there would also be no exteroceptive prediction errors and Rodney’s interoceptive data would be fully explained. However, if Derek was not empathic, then exteroceptive prediction errors would be generated, which could be resolved by biasing perception of Derek’s expression such that it appears more pained (exteroceptive action), a form of ‘emotional projection’, or egocentric bias. Under Bayesian theories of perception, this process would be formalised as the exteroceptive predictions acting as a prior, which when combined with sensory evidence to form the percept (i.e., the posterior), act to cause Rodney’s expression to be perceived by Derek as more pained than the sensory evidence alone would suggest.

While turn-taking in songbirds has been successfully modelled using the predictive processing framework (Friston & Frith, 2015), empirical evidence for interpersonal effects of hierarchical generative models as specified by the predictive processing framework is scarce. Despite plentiful evidence of another’s state impacting that of the self (Blakemore, Bristow, Bird, Frith, & Ward, 2005; Chapon, Perchet, Garcia-Larrea, & Frot, 2019; Heyes, 2011; Lamm, Decety, & Singer, 2011; Liu et al., 2019) and several studies demonstrating that one’s own state can influence inference of another’s state (Edey, Yon, Cook, Dumontheil, & Press, 2017; Pezzulo et al., 2018; Rütgen et al., 2015, 2021; Silani, Lamm, Ruff, & Singer, 2013), these empirical studies have not demonstrated, for example, that the degree to which predictions about one’s own state influences perception of another’s state is determined by their precision (a fundamental tenet of predictive processing). It is this prediction that the present study was designed to test.

In brief, an upcoming interoceptive state (pain) was signalled to participants using a cue which afforded a precise or imprecise prediction as to that interoceptive state (i.e., the magnitude of the pain to be experienced). Participants were asked to judge the intensity of a pained facial expression which was presented visually at the same time as the pain was delivered. Crucially, under predictive processing accounts, exteroceptive and interoceptive ‘hypotheses’ about the world outside the brain are biased by expectations. This can be achieved by increasing the precision of units that encode signals the agent expects to encounter (Friston, 2018; Press & Yon, 2019). Accordingly, it was predicted that more precise expectations about participants’ upcoming pain would lead to more precise interoceptive predictions (see Hoskin et al., 2019). The precision of the interoceptive predictions should determine the precision of associated exteroceptive predictions and therefore (under Bayesian perception accounts) the degree to which those exteroceptive predictions influence perception of the other’s state. Accordingly, it was predicted that precise interoceptive predictions about participants’ own pain states should cause a greater influence of this state on perception of the other – specifically that the receipt of painful electrical stimulation should bias perception of another’s pain state more when accompanied by precise interoceptive predictions, than when accompanied by imprecise interoceptive predictions.

## Method

### Participants

In the absence of available data to conduct power calculations, an opportunity sample was collected in which all participants fulfilling the inclusion criteria who responded to the advertisement over six months of data collection were tested. The final sample was composed of 25 females and 24 males between the ages of 18 and 43 years (*M* = 23.5, *SD* = 5.86). All participants had normal or corrected-to-normal vision, rated the maximum electrical stimulation as at least an 8 out of 10 (details below), were not diagnosed with any neurodevelopmental disorder, nor did they meet the criterion for severe alexithymia (20-item Toronto Alexithymia Scale (TAS-20; Bagby, Parker, & Taylor, 1994) score > 60) as alexithymia has been associated with impaired interoception (Brewer, Cook, & Bird, 2016; mean TAS-20 score 41.8, *SD* = 9.04). Participants did not report taking any prescription medications with stimulant, sedative, or analgesic effects. Participants were also asked to have a full night’s sleep before the experimental session, and to refrain from caffeine consumption on the day of testing. Participants were excluded from analyses if they deviated more than three standard deviations from the group mean on measures of pain rating consistency (two participants) or habituation (two participants). All participants gave written informed consent, and the study was approved by the Central University Research Ethics Committee, University of Oxford. Participants received a small honorarium for their participation.

### Electrical Stimulation and Thresholding

Pain stimuli consisted of 200 *μ*s electrical pulses generated by a Digitimer DS7A Constant Current Stimulator (Digitimer Ltd, Hertfordshire, United Kingdom). Stimuli were controlled by a custom MATLAB script and administered via a bar electrode (two disc electrodes with 9 mm diameter and 30 mm spacing) attached to the underside of the forearm of the non-dominant hand.

Stimulation levels were calibrated for each participant, creating a personalized ‘1’ to ‘10’ scale of pain. A value of ‘1’ corresponded to a minimally painful pin-prick sensation, while ‘10’ was the most painful stimulation participants were willing to receive up to 30 times over the following hour, which did not cause wincing, blinking, or a lapse in focus. Each participant received an ascending series of electrical stimulations, starting at an imperceptible level (1 mA), until they reported first feeling a painful pin-prick sensation. Starting from above this value, a series of stimulations of descending intensity was given until participants reported no longer feeling the pin-prick sensation. The ascending and descending painful thresholds were averaged to give the participant’s ‘1’ value. The intensity level was then further increased until the participant reported reaching ‘10’. Again, starting from above this value, a descending series of stimulations was given until participants reported the intensity dropping below ‘10’ value, and the ‘10’ value was taken as the average of the ascending and descending thresholds. The mean difference between ‘10’ and ‘1’ stimulation intensities was 40.1 mA (*SD* = 22.0). Provisional stimulation levels for values ‘2’ through ‘9’ were calculated as equidistant points between the ‘1’ and ‘10’ values. For each value, the provisional stimulation level was adjusted via further calibration according to participant feedback in increasingly fine intervals until the participant’s subjective rating matched the assigned value.

### Measures of Pain Reporting and Degree of Habituation

Before the main task, in a pre-test phase, participants received each of their 10 individually-calibrated stimulation intensities twice. The order of intensities was random, but held constant across all participants so that any effects of order on pain perception would be equal across participants. Participants were asked to rate each stimulation out of 10, based on the scale used during calibration. From these data, estimates of participant accuracy (correlation between the average of the two pre-test ratings and the actual intensity level) and consistency (correlation between the first and second pre-test rating for each shock level) were calculated. After the main task, in a post-test phase, this procedure was repeated, with each stimulation level being presented only once. Comparison of the pre- and post-test data allowed a measure of habituation to be derived (the mean difference between the post-test and the average of the pre-test ratings across intensity levels) for each participant.

### Emotional Facial Expression Stimuli

Stimuli were images of a female actor displaying happy and pained facial expressions of varying intensities, created by morphing each expression with a neutral expression using Morpheus Photo Morpher (Morpheus Development, Howell, Michigan). Original stimuli were obtained and validated by Simon, Craig, Gosselin, Belin, & Rainville (2008). Morphed images were converted to grey-scale and cropped into an oval shape to occlude hair, neck and peripheral information.

For both pain and happiness, 18 intermediate images between the neutral (0%) and the emotional expression (100%) were initially produced in 5% increments. A pilot study (n = 50) conducted using these images revealed that participants required 10% more happiness in happy morphs than the amount of pain required in pain morphs to judge the facial image as happy/pained, respectively. Therefore, to equalize perceived intensity of the two emotions, the final happy stimuli consisted of five morphs selected from a range of intensities (minimum 35%, maximum 70% intensity) each of which were 10% more intense than the corresponding pained morphs (minimum 25%, maximum 60% intensity; Figure 1). Stimuli were 222 x 293 pixels in size, presented on a grey background in Psychtoolbox (Brainard, 1997) and viewed from a distance of approximately 60 cm. Presentation time was 425 ms.

**Figure 1.** Figure removed from preprint.

### Pain Cues

In order to manipulate the precision of pain predictions, participants were presented with a cue prior to receiving each stimulation that informed them, with high or low precision, whether they were going to receive a high- or a low-pain stimulation. Cues were shown as horizontal bars, signifying the range from minimum (1) to maximum (10) pain, with a shaded region indicating the range of possible intensities of the upcoming stimulation. For low precision cues, this shaded region occupied 50% of the bar, indicating that the pain could be anywhere from minimum to mid-way (for low pain) or mid-way to maximum (for high pain). For high precision cues, 10% of the bar was shaded, centred around 25% (for low pain) and 75% (for high pain).

### Design

The design consisted of three variables manipulated on a within-subjects basis: pain stimulation magnitude (Own-Pain: high or low), precision of pain expectation (Precision: high or low) and the type of expressed emotion (Emotion: pain or happiness) and trials representing the factorial combination of these three factors were presented equally over 120 trials in blocks of 24 trials. Blocks consisted of equal numbers of trials from every combination of experimental factors, presented in a random order. In low precision conditions, each facial image was paired once each with a stimulation of level ‘1’, ‘3’, and ‘5’ (in the low own-pain condition) or a stimulation of level ‘6’, ‘8’ and ‘10’ (in the high own-pain condition). In the high precision conditions, the stimulation given was always ‘3’ in the low own-pain condition and ‘8’ in the high own-pain condition. This ensured that the mean stimulation intensity received was equal across high and low precision conditions (i.e., ‘3’ or ‘8’) and also within each facial image.

### Procedure

After obtaining informed consent, the electrode was attached, the calibration procedure carried out, and the pre-test stimulation ratings obtained. There were six practice trials for the main task, presented in a random order but the conditions of which were fixed to include: each combination of Precision and Own-Pain conditions; the most extreme painful stimulations for low precision conditions (i.e., 1 and 5 for low own-pain and 6 and 10 for high own-pain), to reinforce the idea that low precision cues signal a wide range of potential upcoming pain relative to high precision cues, and the most and least intense facial images, so that participants could be instructed to calibrate their scale for rating the emotions accordingly (i.e., the least and most happy/pained expressions should correspond to ‘1’ and ‘10’ on the scale, respectively).

The structure of each trial of the main task is shown in Figure 2. Participants were presented with the own-pain cue for three seconds before being presented with the facial expression for 425 milliseconds. The electrical stimulation was delivered simultaneously with the presentation of the facial expression. Participants were then asked to judge the intensity of the emotional expression, the intensity of their own pain (both on a scale of 1 to 10), and whether the facial expression was happy or pained. Participants were encouraged to take a break between blocks. After the main task, the post-test rating procedure was carried out.

**Figure 2.**
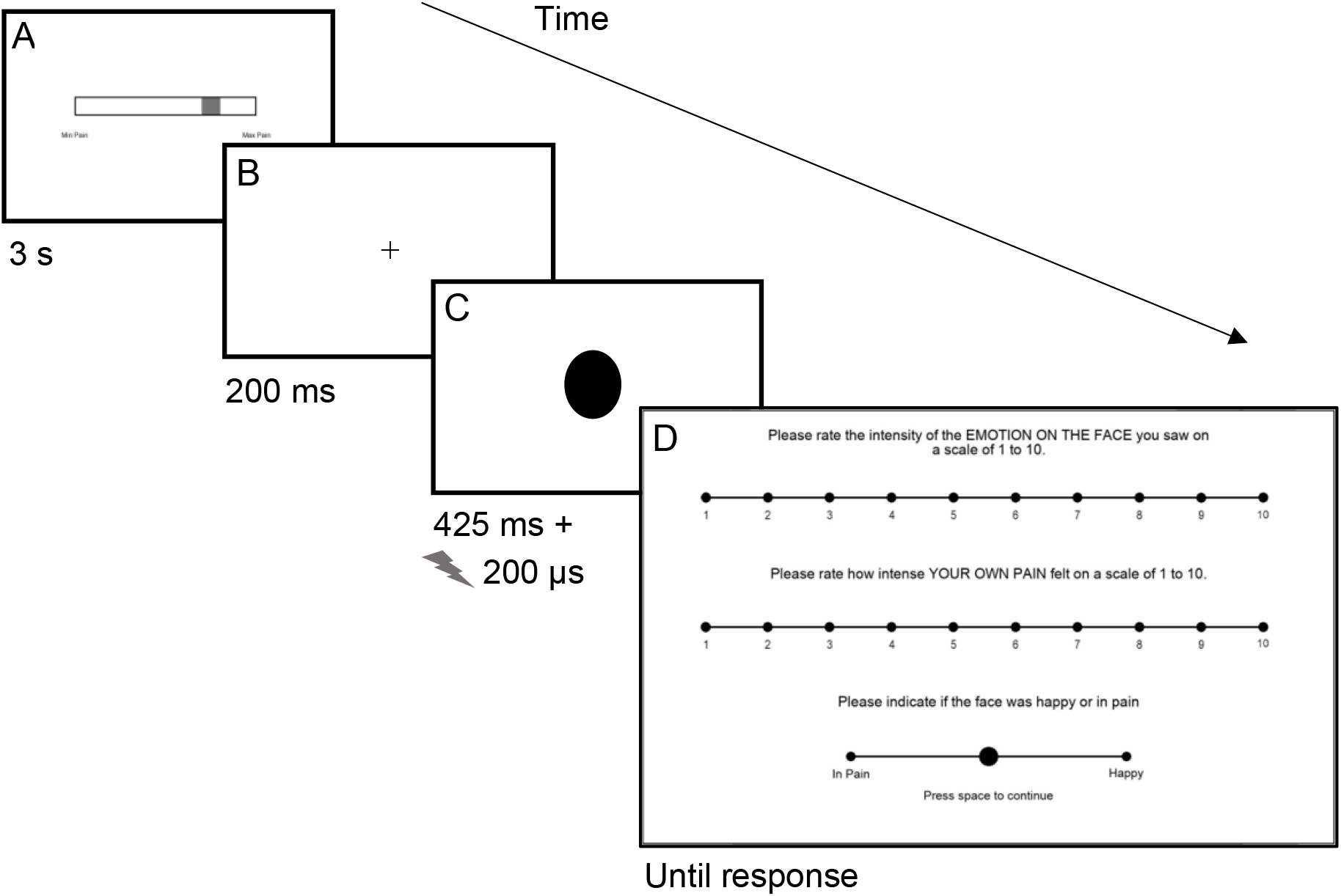
Task Structure. *Note*. Example cue and stimulus shown – these varied across trials as specified under ‘Design’. A) Cue: Indicates the magnitude of the upcoming electrical stimulation (High or Low own-pain) with either a High or Low degree of precision (High Pain, High Precision cue shown); B) ISI; C) Expression stimulus: Either Pained or Happy with concurrent electrical stimulation (facial stimulus removed from image); D) Response Screen: Own pain rating (1-10) + Expression Intensity rating (1-10) + Emotion judgment (pained or happy).

### Results

All statistical analyses were computed in JASP (Jasp Team, Amsterdam, the Netherlands). All tests are two-tailed unless otherwise specified. Bayesian analyses use JASP default priors.

### Pre- and Post-Test Own-Pain Ratings

The mean consistency correlation for own-pain rating (within-participant correlation between the two pre-test ratings) was .87 (*SD* = .09) and the mean accuracy correlation (within-participant correlation between the mean pre-test ratings and the calibrated pain levels) was .95 (*SD* = .03). The mean habituation score was 0.09 (*SD* = 0.64), corresponding to a slight habituation.

### Expression Intensity Ratings

Expression intensity ratings (see Figure 3) were analysed using a 2 (Own-Pain: high vs. low) x 2 (Precision: high vs. low) x 2 (Emotion: pain vs. happiness) repeated measures analysis of variance (ANOVA). As predicted, there was a significant 2-way interaction between Own-Pain and Emotion [*F*(1, 48) = 5.61, *p* = .022, η_p_^2^ = .11], and crucially, a significant 3-way interaction between Own-Pain, Precision and Emotion [*F*(1, 48) = 11.4, *p* = .001, η_p_^2^= .19]. There were also significant main effects of Own-Pain [*F* (1, 48) = 42.6,*p*< .001, η_p_^2^= .47] and Emotion [*F*(1, 48) = 43.0, *p*< .001, η_p_^2^= .47]. All other main effects and interactions were non-significant and not of theoretical relevance [all *F*(1, 48) ≤ 0.47, *p* ≥ .497, η_p_^2^≤ .01].

**Figure 3.**
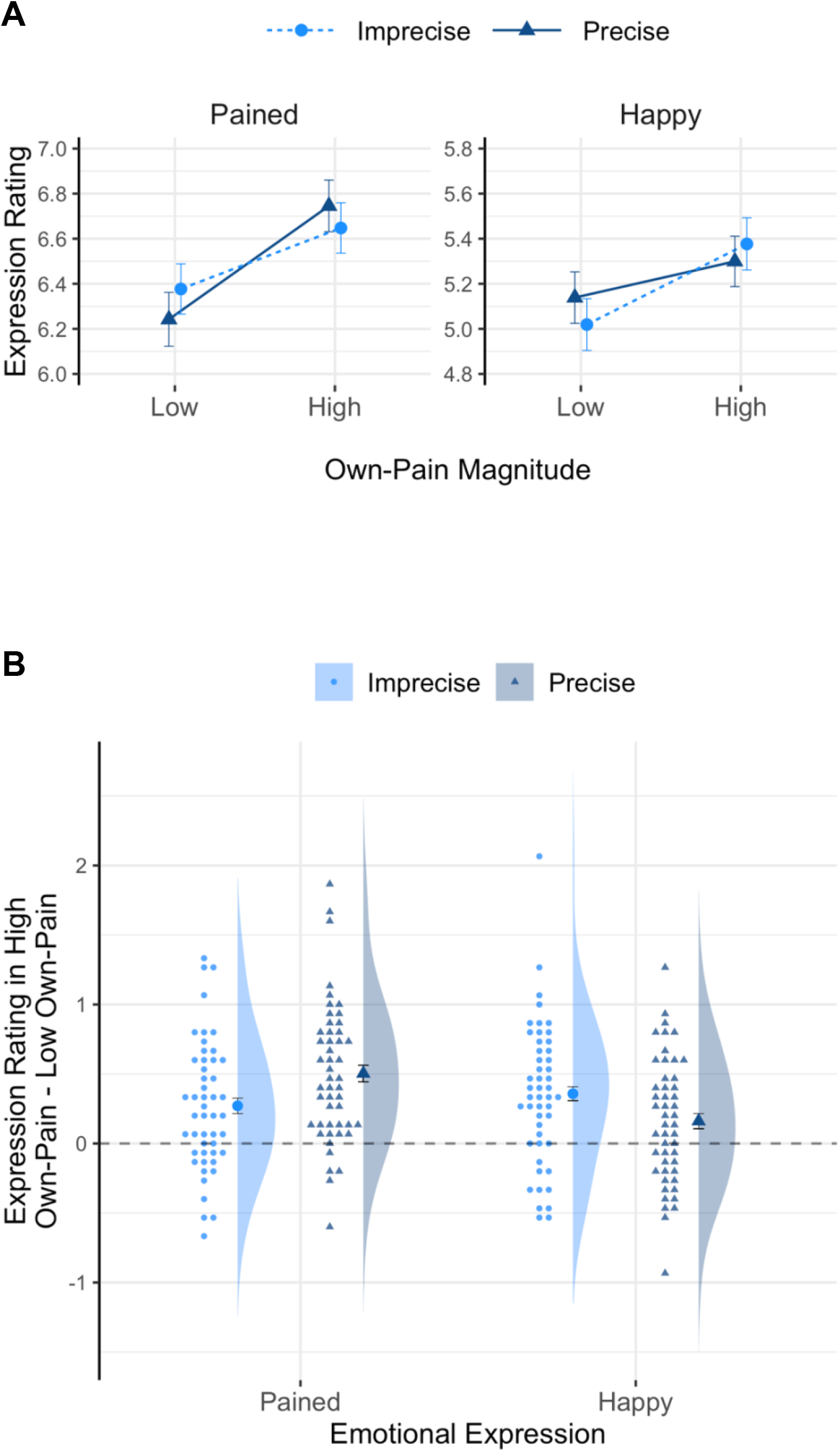
Expression Ratings as a Function of Own-Pain and Precision. *Note*. Panel A: mean rating of expression intensity as a function of own pain magnitude and precision for pained and happy expressions. Panel B: difference in expression rating between High and Low Own-Pain conditions for each combination of precision and emotion. Raincloud plots provide data distributions, mean values and raw data (jittered on the x axis). Error bars show within-subject standard error (Morey, 2008).

To deconstruct the 3-way interaction, two separate 2 x 2 repeated measures ANOVAs were performed for pain and happiness. To investigate these 2-way interactions and the significant 2-way interaction between Own-Pain and Emotion, paired samples t-tests were performed and supplemented by Bayes factors (BF_10_), using the framework proposed by Jeffreys (1961, see also Rouder, Speckman, Sun, Morey, & Iverson, 2009). The Bayes factors reflect how many times more likely the data are under the alternative hypothesis (that there is a difference in expression ratings between the relevant conditions) relative to the null (that there is no difference in expression ratings between the relevant conditions).

The simple main effect of Own-Pain on expression ratings (‘mean difference’ refers to expression ratings in high Own-Pain subtracted from low Own-Pain conditions) was significant for both Emotion conditions, both across and within Precision conditions, but was greater for pained expressions [mean difference =0.39, *SD* = 0.38; *t*(48) = 7.09, *p* < .001, *d* = 1.01; BF_10_ = 2.28×10^6^] than happy expressions [mean difference = 0.26, SD = 0.41; *t*(48) = 4.46, *p*< .001, *d*= 0.64; BF_10_ = 435]. As predicted, and as evidenced by a significant two-way interaction between Own-Pain And Precision (*F*(1, 48) = 7.61, *p* = .008, η_p_^2^ = .14), the effect of Own-Pain on pained expressions was greater in the high precision [mean difference = 0.50, *SD* = 0.50; *t*(48) = 7.05, *p*< .001, *d* = 1.01; BF_10_ = 2.01×10^6^] than the low precision [mean difference = 0.27, *SD* = 0.47; *t*(48) = 4.07,*p*< .001, *d* = 0.58; BF_10_ = 139] condition. Conversely, the simple main effect of Own-Pain on ratings of happy expressions was greater in the low precision [mean difference = 0.36, *SD* = 0.51; *t*(48) = 4.90, *p*< .001, *d* = 0.70; BF_10_ = 1,693] than the high precision [mean difference = 0.16, *SD* = 0.45; *t*(48) = 2.49, *p* = .016, *d* = 0.36; BF_10_ =2.48] condition (see Figure 3), and this interaction between Own-Pain and Precision was significant (*F*(1, 48) = 7.10, *p* = .010, η_p_^2^ = .13).

These results are confirmed by a one-tailed Bayesian paired samples t-test comparing the 2-way interaction effects (computed as the difference in the effect of pain on expression ratings between high and low precision conditions) for happy and pained expressions. A BF_10_ of 41 constitutes strong evidence for the predicted interaction between Pain, Precision and Emotion.

### Confirmatory and Control Analyses

If the effect on expression intensity ratings is as predicted by the predictive processing framework (and Bayesian perception accounts in general), one would expect an effect of cue precision on the variance of own-pain ratings. Precise interoceptive predictions as to the intensity of the upcoming painful stimulation would be combined with sensory evidence to form a precise posterior distribution, leading to lower variance in reported own-pain given the same sensory evidence (i.e., to stimulations of equal intensity). Conversely, imprecise priors would be combined with sensory evidence to form an imprecise posterior distribution, and higher variance in own-pain perception for stimulations of equal intensity (see Hoskin et al., 2019). As a confirmatory analysis therefore, the variance of own-pain ratings was calculated for stimulations at the ‘3’ and ‘8’ levels (the two stimulation intensities shared in the precise and imprecise distributions) for each participant. Variance was calculated after equalising trial numbers in the precise and imprecise conditions by randomly sampling from the precise condition. These intensities were analysed using a one-tailed paired samples t-test which revealed a significant difference in the variance of own-pain ratings, *t*(49) = 2.00, *p* = .026, *d* = 0.29; BF_10_ = 1.88, although note that the Bayes factor provided only anecdotal evidence in favour of the alternative hypothesis (likely due to low power as a consequence of reduced trial numbers).

A control analysis was conducted to ensure that the observed effects were due to the precision of interoceptive cues affecting the precision of exteroceptive predictions (and therefore the degree to which exteroceptive predictions biased perception), rather than being a product of either of two alternative mechanisms. The first alternative is that the precision of interoceptive predictions affected the mean magnitude of experienced pain, with the relationship between experienced pain and expression intensity judgements remaining constant. The second alternative is that the emotional expression may have affected the experienced pain magnitude, since the predictive processing framework predicts bidirectional biasing effects whereby not only can the experience of pain cause an expression to be perceived as more pained to reduce exteroceptive prediction errors, but the sight of a pained expression can cause pain to be experienced as more intense to reduce interoceptive prediction errors. In order to rule out these alternative explanations, the own-pain ratings were therefore analysed using the same 2 (Own-Pain: high vs. low) x 2 (Precision: high vs. low) x 2 (Emotion: pain vs. happiness) repeated measures ANOVA as used to analyse the expression intensity ratings, and supplemented with a Bayesian version of the same test (Rouder, Morey, Speckman, & Province, 2012). Exclusion Bayes factors (BF_excl_) are reported, calculated for ‘matched’ models; these indicate how many times more likely the data are under models that do not include the predictor than under equivalent models with the predictor.

The ANOVA revealed no significant main effect of Precision [*F*(1, 48) = 3.70, *p* = .060, η_p_^2^ = .072; BF_excl_ = 5.30]. While the frequentist ANOVA revealed a main effect of Emotion on experienced pain [*F*(1, 48) = 7.75, *p* = .008, η_p_^2^ = .14] such that own-pain was rated significantly higher when viewing pained faces (*M* = 5.14, *SD* = 0.52) than when viewing happy faces (*M* = 5.06, *SD* = 0.58), a BF_excl_ of 2.74 suggests that the data provide more evidence in favour of the null hypothesis. Neither the 2-way interactions (Precision x Own-Pain: *F*(1, 48) = 0.009, *p* = .926, η_p_^2^ = .0002, BF_excl_ = 6.88; Precision x Emotion: *F*(1, 48) = 0.014, *p* = .907, η_p_^2^ = .0003, BF_excl_ = 6.53; Emotion x Own-Pain: *F*(1, 48) = 0.014, *p* = .907, η_p_^2^ = .0003, BF_excl_ = 6.11), nor the crucial three-way interaction were significant [*F*(1, 48) = 0.78, *p* = .381, η_p_^2^ = .016; BF_excl_ = 10.1). The pattern of significance therefore does not suggest that the effects of either the precision of interoceptive cues or emotional stimulus on experienced own-pain explain the effect of the interoceptive cues on expression intensity ratings. Even if one ignores the pattern of significance and Bayes factors, given that a difference in own-pain ratings of 5 points was necessary to produce a mean difference of 0.32 in expression intensity ratings, it is unlikely that mean differences in own-pain approximately 90 times smaller than that between precision conditions, and 60 times smaller than between emotion conditions, could account for effects on expression intensity ratings.

## Discussion

This study sought to test the hypothesis that the precision of interoceptive predictions regarding one’s own state determine the effect that state has on perception of another’s state. Results supported the hypothesis; precise interoceptive predictions about upcoming pain in the self resulted in that pain having a greater effect on judgement of the intensity of another’s pained expression than imprecise predictions. Furthermore, this effect was specific to pained expressions; the effect of the precision of interoceptive predictions on ratings of the intensity of happy expressions was significantly smaller than that on pained expressions, and in the opposite direction, such that less precise interoceptive predictions were associated with the greatest effect on expression intensity ratings.

Hypotheses as to the effect of precision were based on the description of hierarchical generative models under the predictive processing framework (e.g., Barrett & Simmons, 2015; Demekas, Parr, & Friston, 2020; Ondobaka et al., 2017; Pezzulo, 2014; Pezzulo, Rigoli, & Friston, 2015; Quattrocki & Friston, 2014; Seth, 2013; Seth & Friston, 2016). These models generate multimodal predictions and therefore can link interoceptive, exteroceptive, and proprioceptive states. This property, combined with a developmental environment in which states of the self reliably predict, and are predicted by, states of the other, allow predictions concerning the other to influence the self and vice versa (Bird & Viding, 2014; Heyes & Bird, 2007; Ondobaka et al., 2017; Quattrocki & Friston, 2014; Seth & Friston, 2016). Such models are therefore consistent with the idea that learning resolves the ‘correspondence problem’ (whereby information about the state of another is typically acquired through exteroceptive senses such as vision and audition, while states of the self are typically encoded in interoceptive or proprioceptive codes) inherent in interpersonal influence (Cook, Bird, Catmur, Press, & Heyes, 2014).

In explaining how interpersonal influence arises, one must also explain how such effects can be overcome, or why it is not the case that we compulsively copy others’ actions (echopraxia) or mirror their emotions, and why pairs of individuals do not become locked into such positive feedback loops. Predictive processing models posit that in order to avoid emotional echopraxia when confronted with another’s pain, one must reduce the precision of interoceptive information — in particular, interoceptive predictions or the ensuing prediction errors that would otherwise engage autonomic reflexes to perform the interoceptive action (i.e., induce the state of pain in oneself). With respect to the effect observed here – where the state of the self influences perception of another’s state – one would need to reduce the precision of exteroceptive predictions/prediction errors (Ondobaka et al., 2017; Quattrocki & Friston, 2014; Seth & Friston, 2016). The process of interpersonal matching (whether emotional echopraxia or emotional projection) due to enhancement of predicted consequences followed by later suppression of predicted effects is consistent with a recent model which suggests that predicted events are subject to enhanced processing and then subsequently suppressed (Press, Kok, & Yon, 2020). It is also consistent with models of empathy which suggest that empathy for another’s pained state develops from simple state matching, likely to lead to personal distress in the empathiser, to a situation in which the empathiser distinguishes between their state and that of the other to develop empathic concern or compassion in which their state diverges from that of the other (e.g., Decety & Lamm, 2006; Quattrocki & Friston, 2014).

In addition to an effect of own-pain on the perception of another’s pain, there was a (smaller) effect of own-pain on the perception of happiness. Possibly, high arousal states in the self enhance perception of all other emotions (though this is contrary to the results of Pezzulo et al. (2018)), or specifically of emotions with a similar degree of arousal as one’s own state, if emotions are conceptualised within the circumplex model (whereby all emotions can be characterised within a two-dimensional space with dimensions of valence and arousal; Russell, 1980). Empirical evidence for an analogous idea regarding valence is provided by Antico, Cataldo, & Corradi-Dell’Acqua (2019), who showed that a pained state enhances perception not only of pain but also, to a lesser degree, disgust (also negative valence). In contrast to the effect of self-pain on perception of pain, however, the effect of self-pain on perception of happiness was reduced, not enhanced, by precise interoceptive predictions. This result suggests that more precise interoceptive predictions relating to one’s own pained state result in more precise exteroceptive predictions, enhancing effects on the perception of pain and reducing effects on the perception of happiness.

The ability of interoceptive predictions to bias exteroceptive perception, as shown here, is consistent with accounts which suggest that interoception biases attentional, sensory and behavioural responses to stimuli that are homeostatically relevant (e.g., Barrett & Simmons, 2015). As argued by Seth and Friston (2016), the predictive processing framework, in particular active inference, highlights the relevance of predictive models to the regulation (not just prediction) of causes of sensory data. Due to their influence on our own states, the states of others are homeostatically relevant, and thus a target for regulation by predictive models. Consequently, it has been suggested that atypical predictive processing may lead to atypical sociocognitive ability, with Autism Spectrum Disorder most frequently cited as a condition where this may be the case (Brock, 2012; Coll, Whelan, Catmur, & Bird, 2020; Pellicano & Burr, 2012; Quattrocki & Friston, 2014; but see Brewer, Happé, Cook, & Bird, 2015).

It is not only atypical predictive processing which may result in a failure to perceive, predict and/or regulate the states of others. The generative models giving rise to multimodal predictions concerning the state of the self and others are a product of experience, and therefore depend on sufficient caregiver-child interaction, and may be subject to individual, familial, and cultural variance (Conway, Catmur, & Bird, 2019; Demekas et al., 2020; Happé & Frith, 1996; Jack, Caldara, & Schyns, 2012; Russell, 1991; Smith, Parr, & Friston, 2019). Such variance may mean that predictive models are appropriate for some individuals, or groups, but not others, and that therefore social interaction and communication with members of groups characterised by similar generative models as the self may well be easier than with those with different generative models (Schuster et al., 2021; Edey et al., 2017; Friston & Frith, 2015; Keating & Cook, 2020; Seth & Friston, 2016).

## Data Availability

Data are available at https://osf.io/4p5ur/.

